# Genome-reconstruction for eukaryotes from complex natural microbial communities

**DOI:** 10.1101/171355

**Authors:** Patrick T. West, Alexander J. Probst, Igor V. Grigoriev, Brian C. Thomas, Jillian F. Banfield

## Abstract

Microbial eukaryotes are integral components of natural microbial communities and their inclusion is critical for many ecosystem studies yet the majority of published metagenome analyses ignore eukaryotes. In order to include eukaryotes in environmental studies we propose a method to recover eukaryotic genomes from complex metagenomic samples. A key step for genome recovery is separation of eukaryotic and prokaryotic fragments. We developed a kmer-based strategy, EukRep, for eukaryotic sequence identification and applied it to environmental samples to show that it enables genome recovery, genome completeness evaluation and prediction of metabolic potential. We used this approach to test the effect of addition of organic carbon on a geyser-associated microbial community and detected a substantial change of the community metabolism, with selection against almost all candidate phyla bacteria and archaea and for eukaryotes. Near complete genomes were reconstructed for three fungi placed within the eurotiomycetes and an arthropod. While carbon fixation and sulfur oxidation were important functions in the geyser community prior to carbon addition, the organic carbon impacted community showed enrichment for secreted proteases, secreted lipases, cellulose targeting CAZymes, and methanol oxidation. We demonstrate the broader utility of EukRep by reconstructing and evaluating relatively high quality fungal, protist, and rotifer genomes from complex environmental samples. This approach opens the way for cultivation-independent analyses of whole microbial communities.

## Introduction

Microbial eukaryotes are important contributors to ecosystem function. Gene surveys or DNA “barcoding” are frequently used to identify eukaryotes in microbial communities and have demonstrated the breadth of eukaryotic diversity (Pawlowski et al. 2012). However, these approaches can only detect species and are unable to provide information about metabolism or lifestyle in the absence of sequenced genomes. The majority of fully sequenced eukaryotic genomes are from cultured organisms. Lack of access to cultures for a wide diversity of protists and some fungi detected in gene surveys has resulted in major gaps in eukaryotic reference genome databases (Caron et al. 2008, Pawlowski et al. 2012). Single cell genomics holds promise for sequencing uncultured eukaryotes and has generated partial genomes for some (Cuvelier et al., 2010; Yoon et al., 2011; Monier et al., 2012; Vaulot et al., 2012, Roy et al. 2014, Mangot et al. 2017). However, multiple displacement amplification limits the completeness of single cell genomes (Woyke et al., 2010). Alternatively, metagenomic sequencing reads from environmental samples are mapped against reference genomes to detect organisms and constrain metabolisms, but this approach is restricted to study of organisms with sequenced relatives.

Many current studies of natural ecosystems and animal- or plant-associated microbiomes use an untargeted shotgun sequencing approach. When the DNA sequences are assembled, tens of thousands of genome fragments may be generated, some of which derive from eukaryotes. Exceedingly few metagenomic studies have systematically identified such fragments as eukaryotic, although some genomes for microbial eukaryotes have been reconstructed (Sharon et al. 2013, Quandt et al. 2015, Kantor et al. 2015, Mosier et al 2016, Raveh-Sadka et al. 2016, Kantor et al. 2017). In almost all cases, these genomes were recovered from relatively low diversity communities where binning of genomes is typically less challenging than in complex environments. Here we applied a new kmer-based approach for identification of assembled eukaryotic sequences in datasets from diverse environmental samples. Identification of eukaryotic genome fragments enabled their assignment to draft genomes and improvement of the quality of gene predictions. Predicted genes on assembled metagenomic contigs provide critical inputs for further binning decisions that incorporate phylogenetic profiles as well as classification of the reconstructed genomes and assessment of their completeness. Our analyses focused on biologically diverse environmental samples, many of which came from groundwater. In addition, we investigated previously published metagenomes from infant fecal samples and a bioreactor community used to break down thiocyanate. Because the approach works regardless of a pre-determined phylogenetic affiliation, it is now possible to reconstruct genomes for higher eukaryotes as well as fungi and protists from complex environmental samples.

## Results

### Crystal Geyser Community Structure

The deep subsurface microbial community at Crystal Geyser (Utah, USA) has been well characterized as being dominated by chemolithoautotrophic bacteria and archaea, including many organisms from candidate phyla (CP) (Probst et al. 2014, Emerson et al. 2015, Probst et al. 2016). It is our current understanding that a wide diversity of novel bacteria and archaea are brought to the surface by geyser eruptions (Probst et al. 2017 in revision). Such deep sedimentary environments are unlikely to have high organic carbon compound availability. Thus, we hypothesized that organic carbon addition to this system would profoundly shift the community composition by selecting against the novel geyser microorganisms and enriching for better known heterotrophs. To test this prediction, we analyzed a sample of wood that was added to the shallow geyser and had decayed in the groundwater conduit (hereafter referred to as CG_WC). This sample, as well as a wood-free sample (CG_bulk) that was collected the day before CG_WC, were subjected to metagenomic analysis. We identified 124 and 316 distinct strains in the CG_WC and CG_bulk samples respectively. The CG_WC sample contained abundant eukaryotic sequences (Figure 1A) that were not present in the surrounding geyser water (Figure 1B). Twelve strains were present in both samples (Figure 1C), including the archaeon *Candidatus* “Altiarchaeum hamiconexum” (Probst et al. 2014), which dominated the CG_bulk sample. A phylum-level comparison of the microbial communities is presented in Figure 1D. The presence of decaying wood strongly enriched for Actinobacteria and Proteobacteria, as well as eukaryotes such as Ascomycota, Basidiomycota and an organism classified as part of the arthropoda. A low abundance alga from the class bacillariophyta was detected in both samples.

**Figure 1.**
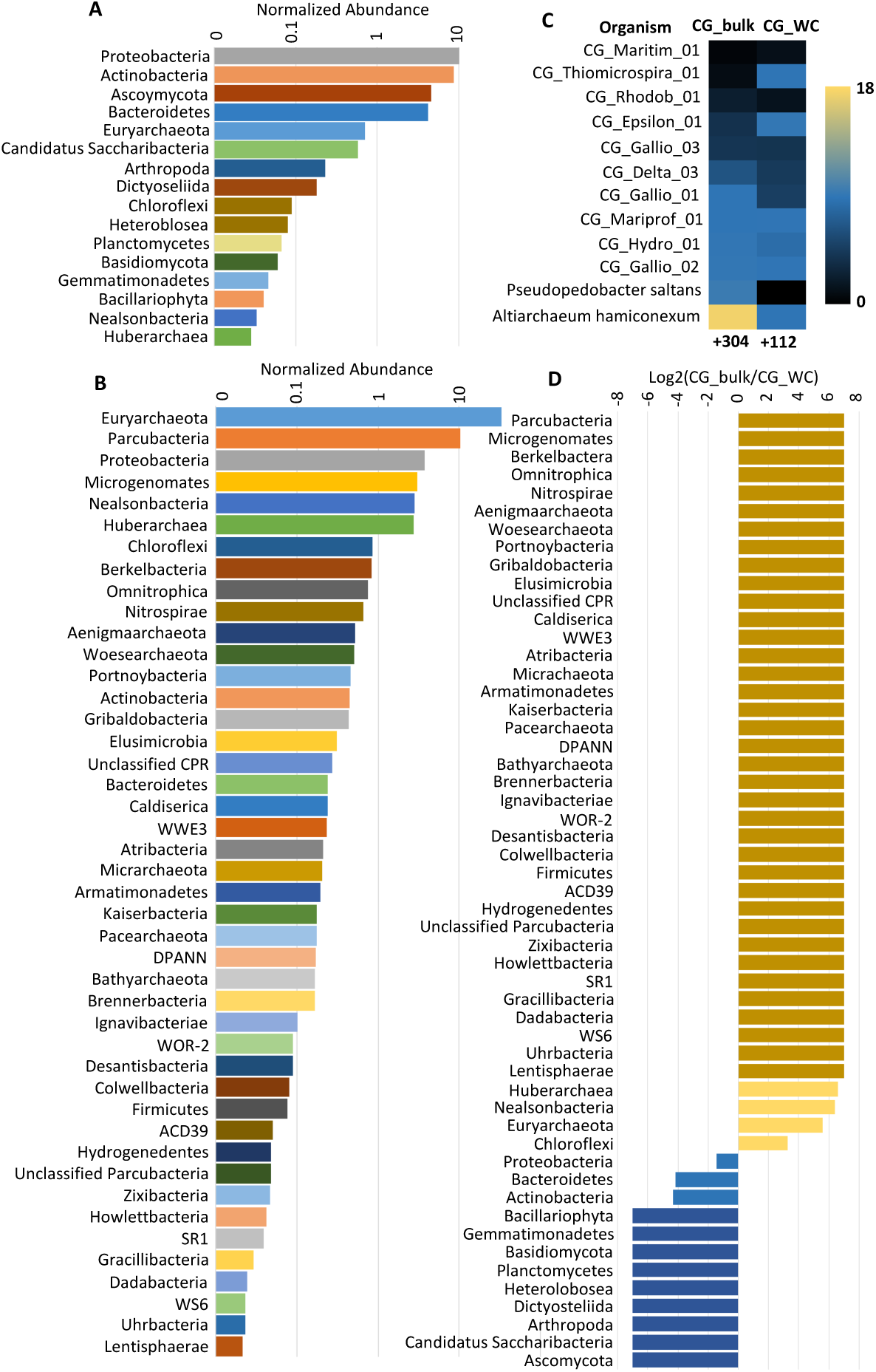
Comparison of CG_WC and CG_bulk community composition. The relative abundances of taxonomic groups in CG_WC (A) and CG_bulk (B) are depicted. Abundance was determined based on the average coverage depth of the scaffolds containing annotated ribosomal protein S3 (rpS3) genes. Abundances were normalized for comparison across samples by multiplying the average coverage depth by the sample read count and read length. (C) Normalized coverage of rpS3 containing scaffolds of strains common to both samples. The number of additional strains detected in each sample are listed below the respective sample heatmap. (D) Log2 ratio of normalized coverage of taxonomic groups from A and B. Taxonomic groups identified in only one sample are indicated by the darker yellow and blue bars.

As predicted, the CG_WC sample contains very few CP bacteria and archaea, with the notable exception of three members of Saccharibacteria (TM7). Two Saccharibacteria genomes were >90% complete and one 1.01 Mbp genome was circularized and curated to completion. To evaluate for the accuracy of the complete genome we ruled out the presence of repeat sequences that could have confounded the assembly and carefully checked the consistency of paired reads mapped across the entire genome (**Supplementary data 1**). The cumulative GC skew was used to identify the origin and terminus of replication. Although the skew has generally the expected form (consistent with genome accuracy), the origin defined based on GC skew was offset from the dnaA gene by ~46 kbp (**Supplementary Figure S1A**). Interestingly, short repeat sequences often associated with the origin were absent both from the predicted origin and the region encoding dnaA, although they were identified close to the origin for another candidate phyla radiation bacterium (Anantharaman et al. 2016). We identified the origin region for a previously reported complete Saccharibacteria RAAC3_TM7 genome using cumulative GC skew and showed that repeats were not present in this genome either, and that the predicted origin is 7.6 kb from the dnaA gene.

### EukRep tested on reference datasets

Typically, only prokaryotic gene prediction is performed on metagenomic samples, as these are the only algorithms specifically designed for this application (e.g., MetaProdigal, Hyatt et al. 2012). For samples containing both prokaryotic and eukaryotic DNA, such as CG_WC, obtaining high quality gene predictions for eukaryotes is complicated by the fact that distinct gene prediction tools are used for prokaryotic vs. eukaryotic sequences due to differences in gene structure. Specifically, eukaryote genomes have more complex promoter regions, regulatory signals, and genes spliced into introns and exons, variable between species. For this reason, it is not surprising that we found that prokaryotic gene predictors underperform when used on eukaryotic sequences (Supplementary **Figure 2**). To address this issue and obtain high quality eukaryotic gene predictions from metagenomes, we present EukRep, a classifier that utilizes kmer composition of assembled sequences to identify eukaryotic genome fragments prior to gene prediction (Figure 2). When previously used to taxonomically classify metagenomic sequences, machine learning algorithms have shown promise, but their success was limited when samples contained many different species (Vervier et al. 2016). We hypothesized a supervised classification method could be applied to accurately classify sequences at the domain level for gene prediction purposes, avoiding complications from having a large number of taxonomic categories.

**Figure 2.**
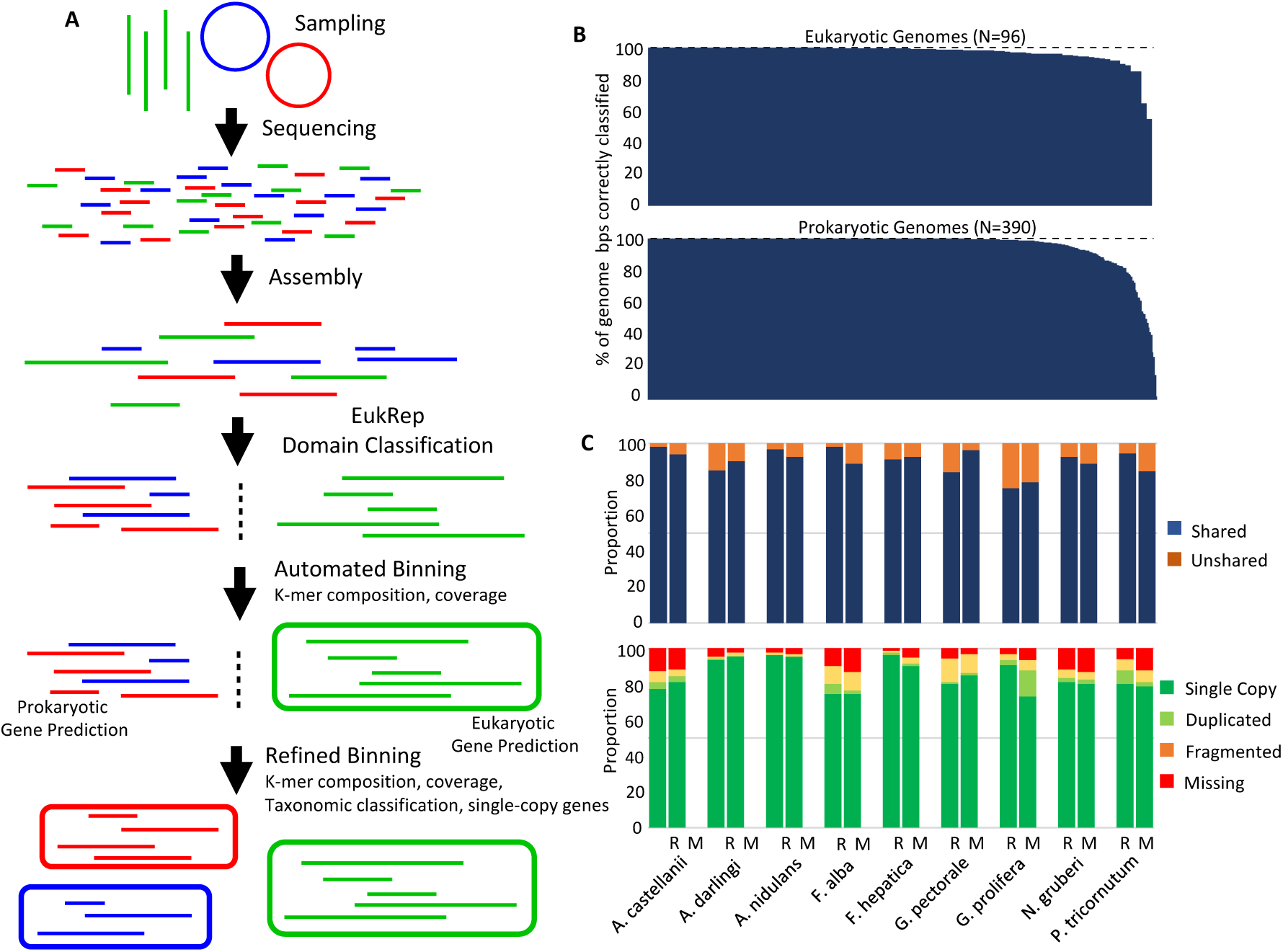
Identification of scaffolds for eukaryotic gene prediction with EukRep. (A)Schematic of the analysis pipeline used to identify and bin both eukaryotic and prokaryotic genomes within this paper. (B) Accuracy of EukRep domain prediction on a per-genome level for both eukaryotes and prokaryotes. Percent of genome correctly classified is defined as the percent of base pairs within a given genome predicted to belong to the genome’s known domain. Each bar represents the percent of a single genome that was classified correctly. Genomes used for training and testing of EukRep are listed in Supplemental Tables S1 and S2 (C) Gene predictions for nine diverse eukaryotic organisms including fungi, a Metazoa, a Stramenopile, an Archaeplastida, and a Rhizaria. Columns labeled “R” refer to reference gene sets whereas M columns refer to gene sets predicted without transcript or close homology evidence. The top panel displays the proportion of total genes either overlapping (shared) or not overlapping (unshared) a gene model from the other respective gene set for a given genome. The bottom panel is an analysis of presence or absence of single copy genes in each gene set as determined by BUSCO using the eukaryota_odb9 lineage set.

The EukRep model was trained using a diverse reference set of bacterial, archaeal, opisthokonta, and protist genomes (3.40 Gbps of sequence; Supplemental Table S1). The kmer frequencies were calculated for each 5-kb interval, resulting in 581,376 individual instances that were used to train a linear-SVM (scikit-learn, Pedregosa et al. 2011). We found that 5-mer frequencies represented the best compromise between speed and accuracy for classifying eukaryotic scaffolds and that sequences can be classified with high accuracy at lengths of 3kb or greater (**Supplemental Figure S3**). Using a validation set of 486 independent genomes (**Supplemental Table S2**) to test EukRep we found that the classifier was able to accurately predict the domain of 97.5% of total tested eukaryotic sequence length and 98.0% of prokaryotic sequence length.

We examined classifier accuracy on a per-genome basis to test whether the classifier performance varied for organisms of widely different types (Figure 2B). This metric differs from that reported above because it refers to the accuracy of classifying individual artificially fragmented genomes rather than overall accuracy on all scaffolds tested from every genome. 94% of tested eukaryotic genomes were classified with >90% accuracy whereas 88% of tested prokaryotic genomes were classified with >90% accuracy. In a small number of prokaryotic genomes more than half of the contigs were misclassified as eukaryotic. Notably, all of these were small genomes of organisms inferred to be parasites or symbionts. However, almost all of the sequences composing the eukaryotic genomes tested were correctly classified, indicating this method can successfully identify scaffolds whose analysis would benefit from a eukaryotic gene prediction algorithm.

### Testing eukaryotic gene predictions on reference genomes

Eukaryotic gene prediction algorithms rely on a combination of transcriptomic evidence or protein similarity (AUGUSTUS, Stanke et al. 2006, SNAP, Korf 2011) and sequence signatures (GeneMark-ES, Ter-Hovhannisyan et al. 2008) to make predictions. Given the frequent lack of sequenced close relatives to organisms identified in metagenomes and the lack of transcript data in many metagenomic studies, we tested how well eukaryotic gene predictors function in a diversity of eukaryotic genomes without transcriptomic evidence or homology evidence from close relatives. We applied the MAKER pipeline (Holt and Yandell, 2011) with GeneMark-ES in self-training mode along with AUGUSTUS trained using BUSCO (Simão et al. 2015) to nine diverse eukaryotic genomes obtained from JGI’s portal (Grigoriev et al. 2011) and NCBI’s genome database (NCBI Resource Coordinates 2017) (Figure 2C). The proteomes of *C*. *reinhardtti* (Merchant et al. 2007), *N*. *crassa* (Galagan et al. 2003), and *R*. *filosa* (Glöckner et al. 2014) were also used as homology evidence. In each case, MAKER-derived gene predictions were compared to reference gene predictions that incorporate transcriptomic evidence. The majority of the gene predictions identified without transcriptomic evidence were supported by reference gene predictions (78-98%) and the majority of reference gene predictions overlapped a MAKER-derived gene prediction (75-98%). Estimated completeness of the predicted gene sets was measured by using BUSCO (Simão et al. 2015) to search for 303 eukaryotic single copy orthologous genes within the predicted gene sets. The number of single copy, duplicated, fragmented, and missing genes showed minimal differences with and without transcriptomic evidence (Figure 2C). These results show the pipeline we assembled for eukaryotic gene prediction, even without transcriptomic evidence, is capable of detecting near complete gene sets similar to those from reference genomes, with exception of untranslated regions and alternative splicing patterns.

### Analysis of newly reconstructed Eukaryotic Genomes

After benchmarking EukRep on reference datasets, the algorithm was applied to the CG_WC sample. 214.8 Mbps of scaffold sequence was classified as eukaryotic. Because eukaryotic gene predictors are designed to be trained and run on a single genome at a time, CONCOCT (Alneberg et al. 2014), an automated binning algorithm, was applied to the identified eukaryotic scaffolds to generate two preliminary eukaryote genomes. In this way, GeneMark-ES and AUGUSTUS gene prediction could be performed, as described above, on each bin individually as if running on a single genome.

The availability of relatively confident gene predictions for eukaryotic contigs enabled re-evaluation of genome completeness based on the presence or absence of 303 eukaryotic single-copy genes as identified by BUSCO (Table 1, Figure 3). An obvious finding was that one of the CONCOCT bins was a megabin. Using information about single-copy gene inventories, along with tetranucleotide frequencies, coverage and GC content, we assigned the eukaryotic scaffolds into four genome bins. Blasting gene predictions against UniProt identified three of the bins as likely fungi and a fourth as a likely metazoan. Gene prediction was redone on the new fungal bins with GeneMark-ES in self-training mode and AUGUSTUS trained with BUSCO. The bins ranged in size from 24.5 Mbps to 99.0 Mbps and encoded between 8947 and 18440 genes. BUSCO single-copy orthologous gene analysis showed all four bins were relatively complete individual genomes based on gene content, with the lowest containing 243/303 (80%) and the highest containing 288/303 (95%) single-copy orthologous genes (Table 1, Figure 3). Some genes expected to be in single copy were duplicated, as is often found with BUSCO analysis of complete genomes. Interestingly, the assembly quality of one bin, WC_Fungi_A, appeared to be quite high, with 50% of its sequences contained in scaffolds longer than 599 kb. We reduced potential contamination of eukaryotic bins with prokaryotic sequence by blasting predicted proteins against Uniprot and removing scaffolds with the majority of best hits belonging to prokaryotic genes.

**Table 1.** Summary of binned eukaryotic genome quality, contamination, and completeness. Eukaryotic genomes identified from CG_WC, infant fecal-derived samples, and thiocyanate reactor samples are listed. Genome completeness is defined as the percent of BUSCO single-copy orthologous genes that were present either in a single copy or duplicated.

| System | Sample | Name | Size (bp) | # genes | # scaffolds | N50 | Completeness (%) |
| --- | --- | --- | --- | --- | --- | --- | --- |
| Crystal Geyser | CG_WC | CG_Fungi_A | 24984438 | 8947 | 80 | 599568 | 95 |
| Crystal Geyser | CG_WC | CG_Fungi_B | 38989301 | 15602 | 4724 | 47173 | 90 |
| Crystal Geyser | CG_WC | CG_Fungi_C | 24500285 | 9955 | 3654 | 16121 | 80 |
| Crystal Geyser | CG_WC | CG_Arthropod | 99046889 | 18440 | 8889 | 17347 | 92 |
| Crystal Geyser | CG_4_10_14_3.00 | CG_Stremenopile | 31157668 | 13749 | 3424 | 11782 | 77 |
| Infant Gut | 182_001 | Infant_A_Fungi_A | 11921609 | 5160 | 265 | 66296 | 94 |
| Infant Gut | 182_002 | Infant_A_Fungi_B | 11426193 | 4999 | 334 | 45092 | 92 |
| Infant Gut | N3_182_000G1 | Infant_A_Fungi_C | 11895925 | 5158 | 262 | 66272 | 94 |
| Infant Gut | N1_023_000G1 | Infant_B_Fungi_A | 12280001 | 5293 | 1002 | 17208 | 88 |
| Infant Gut | b023-d007 | Infant_B_Fungi_B | 12603413 | 5320 | 846 | 20788 | 89 |
| Infant Gut | SP_CRL_000G1 | Infant_C_Fungi_B | 12594614 | 5309 | 885 | 22402 | 87 |
| Thiocyanite Reactor | SCNpilot_cont_1000_bf | Rotifer_A | 32149948 | 16252 | 5055 | 6857 | 69 |
| Thiocyanite Reactor | SCNpilot_cont_1000_p | Rotifer_B | 40079970 | 17172 | 1961 | 25209 | 83 |
| Thiocyanite Reactor | SCNpilot_cont_500_p | Rotifer_C | 46690091 | 19593 | 1702 | 39424 | 87 |
| Thiocyanite Reactor | SCNpilot_cont_750_p | Rotifer_D | 45084683 | 20599 | 4908 | 11284 | 87 |
| Thiocyanite Reactor | SCNpilot_expt_500_p | Rotifer_E | 53918756 | 23992 | 4626 | 15939 | 89 |
| Thiocyanite Reactor | SCNpilot_expt_500_bf | Rotifer_F | 52830794 | 23942 | 5156 | 13016 | 90 |
| Thiocyanite Reactor | SCN_pilot_cont_500_bf | Rotifer_G | 59551575 | 24973 | 3237 | 32744 | 91 |
| Thiocyanite Reactor | SCNpilot_expt_500_p | SCNpilot_expt_500_p | 22299472 | 10815 | 3636 | 6549 | 69 |
| Thiocyanite Reactor | cn_treated | cn_treated | 32902255 | 8342 | 559 | 77947 | 85 |

**Figure 3.**
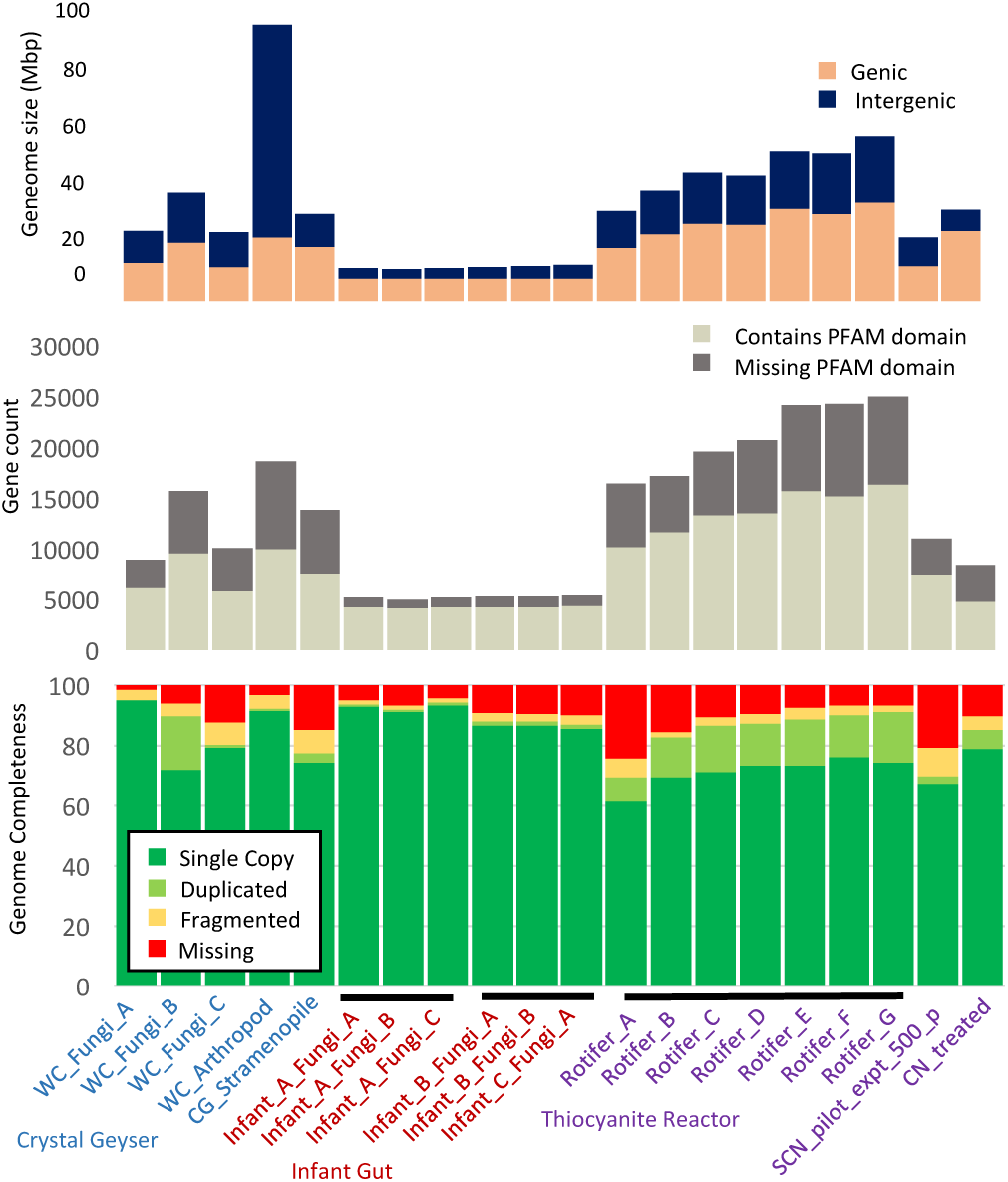
Overview of binned eukaryotic genomes. Genomes that share greater than 99% average nucleotide identity (ANI) are indicated by black bars. ANI comparisons are shown in more detail in Supplemental Figure S3. Genic regions refer to sequence located within predicted gene models whereas intergenic refers to all other sequence. Genes containing a PFAM domain were identified with PfamScan (Mistry et al. 2007). Genome completeness is measured as the percent of 303 eukaryotic single-copy orthologous genes found within a genome in a particular form with BUSCO.

A phylogenetic tree constructed from a set of 16 predicted, aligned, and concatenated ribosomal proteins (Hug et al. 2014) placed three of the bins within the fungal class Eurotiomycetes (Figure 4). Each of these three bins ranged in size from 24.6 Mbps to 39.2 Mbps and in gene count from 8,963 genes to 15,756 genes, within the range observed in previously sequenced Ascomycete fungi. The closest sequenced relative to all three bins was *Phaeomoniella chlamydospora*, a fungal plant pathogen known for causing Esca disease complex in grapevines (Morales-Cruz et al. 2015). The fourth bin, 99.7 Mbps in length and estimated to be 92% complete, was placed within the Arthropoda (Figure 4). Its closest, although distant, sequenced relative is *Orchesella cincta* (Faddeeva-Vakhrusheva et al. 2016). *Orchesella cincta* is a member of the hexapod subclass Collembola (springtails), a diverse group basal to insects known primarily to be detritivorous inhabitants of soil. Although ribosomal protein S3 (rpS3) sequences belonging to Dictyosteliida, Heterolobosea, and Basidiomycota were detected there were no genomes reconstructed for these organisms, likely due to low abundance or genome fragmentation.

**Figure 4.**
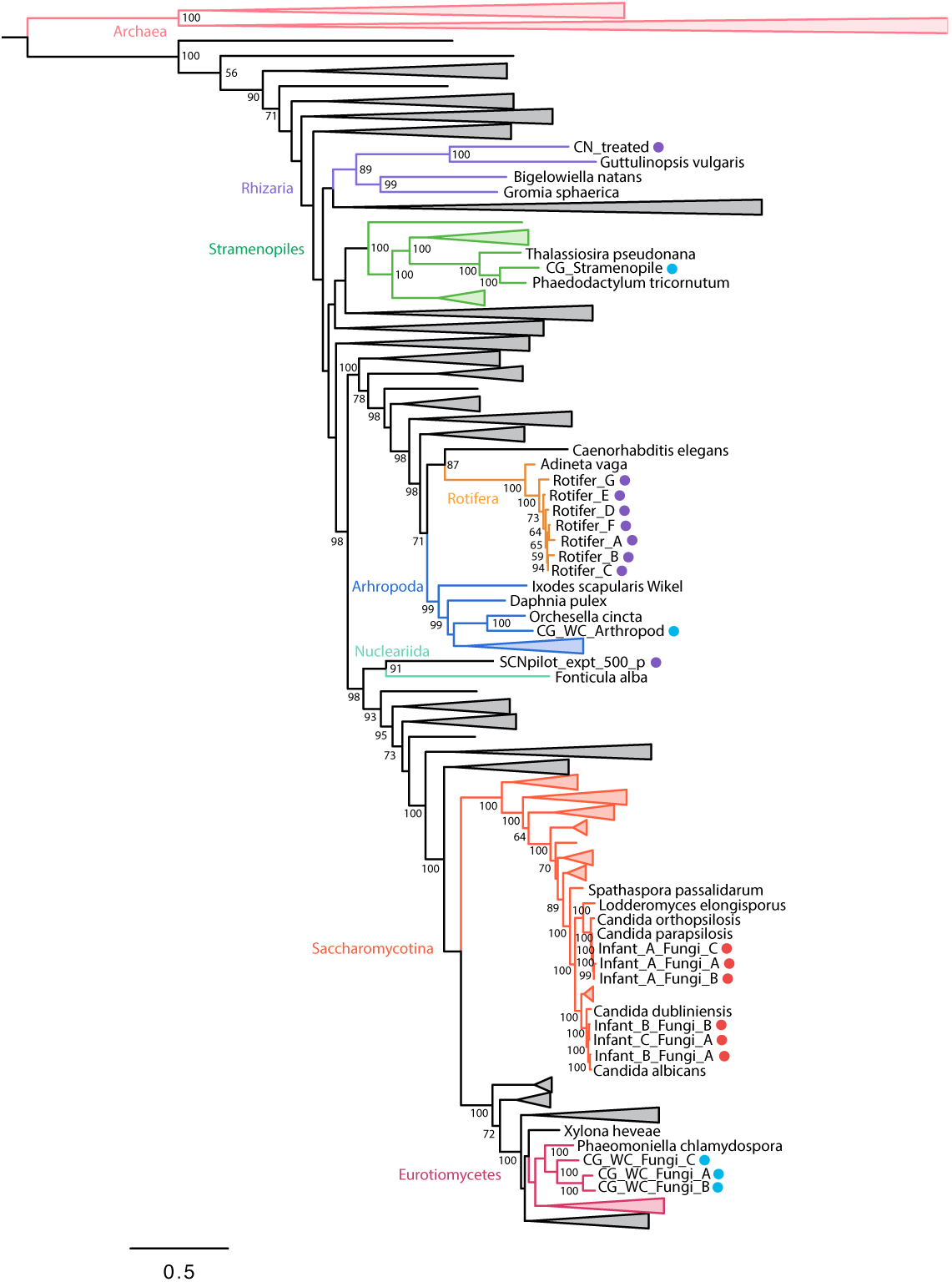
Phylogenetic placement of binned eukaryotic genomes with maximum likelihood analysis of 16 concatenated Ribosomal Protein alignments. Genomes from Crystal Geyser, infant-derived fecal samples, and thiocyanate reactor samples are identified with blue, red, and purple circles respectively. Branches with greater than 50% bootstrap support are labeled with their bootstrap support. Reference ribosomal proteins were obtained from Hug et al. 2016, JGI (Grigoriev et al. 2011), and NCBI (NCBI Resource Coordinators, 2017).

### Whole community analysis, including eukaryotes

To test whether the presence of organic carbon within the CG_WC sample would enrich for heterotrophic metabolic pathways (and against members of chemolithoautotrophic communities typically associated with the Crystal Geyser community), we searched the CG_WC and CG_bulk samples using HMMs for CAZymes grouped by substrate (Cantarel et al. 2009), lipase HMMs from the Lipase Engineering Database (Fischer and Pleiss 2003), and a protease blast database from MEROPS (Rawlings et al. 2016). Predicted proteases and lipases were filtered to specifically identify putative excreted proteases and lipases by searching for proteins with secretion signals identified with SignalP (Petersen et al. 2011) and one or less transmembrane domains with TMHMM (Krogh et al, 2001).

Pathways previously described as dominant within the Crystal Geyser such as the Wood Ljungdahl carbon fixation pathway and Ni-Fe hydrogenases were depleted in CG_WC as compared to CG_bulk. Instead, CAZymes targeting cellulose, hemicellulose, pectin, starch, and other polysaccharides were enriched in CG_WC, indicating an increased capacity for degradation of complex carbohydrates (Figure 5). A strong enrichment for excreted lipases and proteases was also detected, further indicative of an in increase in the amount of heterotrophic metabolisms (Figure 5). Interestingly, CG_WC also had a strong enrichment for methanol oxidation.

**Figure 5.**
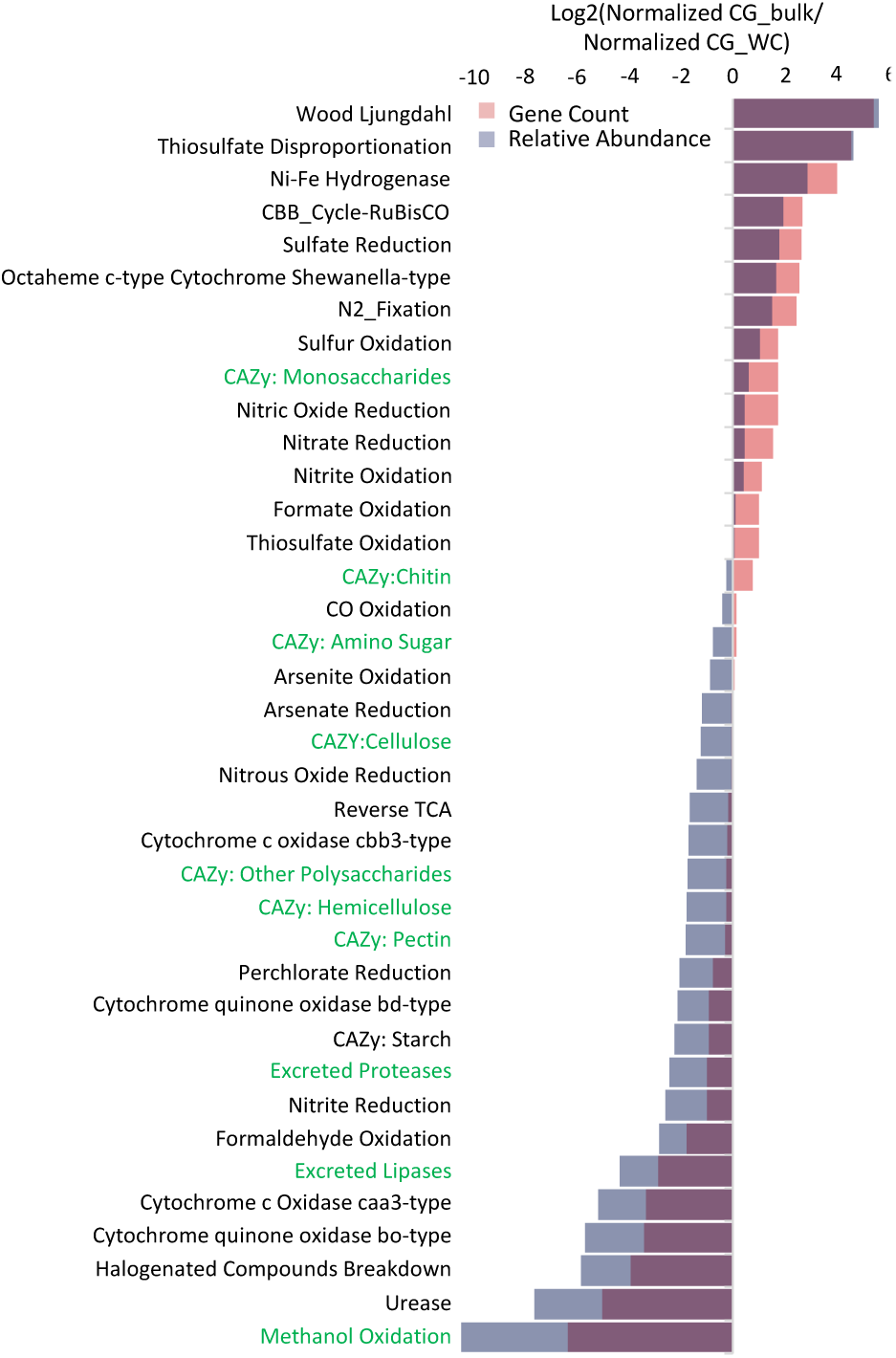
Comparison of CGWC and CGbulk metabolic capacity. Log2 ratio of all annotated genes found within the CG_bulk sample against annotated genes found in the CG_WC sample. Annotated genes were grouped into categories based upon scores with a custom set of metabolic pathway marker HMMs (Anantharaman et al. 2016), CAZYme HMMs (Cantarel et al. 2009), and protease and lipase HMMs from MEROPs and Lipase Engineering Database respectively. Putative proteases and lipases were also filtered to only those containing a secretion signal and one or less transmembrane domains (see methods). Gene count (red) is the ratio of total number of genes in each category for each sample normalized by the total number of genes found in the sample. Relative abundance (blue) is the ratio of average read coverage depth of the contig containing a given annotated gene in each category normalized by the sample read count multiplied by read length

The four binned eukaryotic genomes contributed substantially to the putative heterotrophic categories (**Supplementary Table S3**). Fungi are known to exhibit different CAZyme profiles based upon their lifestyle (Ohm et al. 2012, Kim et al. 2016). An analysis of the CAZyme profiles of the three fungal bins focused on plant cell wall targeting CAZymes supports the role of these fungi as possible plant pathogens or saprotrophs (**Supplementary Table S4**) (Floudas et al 2012, Ohm et al. 2012, Kim et al. 2016). A profile of CAZymes found within the Arthropoda bin revealed a large number of chitin targeting CAZymes (**Supplementary Table S3**).

### Testing EukRep in recovery of Eukaryote genomes from other ecosystems

To test the broader application of EukRep, we applied the method to infant fecal samples and thiocyanate reactor samples in which eukaryotes had previously been identified (Sharon et al. 2013, Raveh-Sadka et al. 2015, Kantor et al. 2015, Raveh-Sadka et al. 2016, Kantor et al. 2017). By using EukRep, we were able to quickly and systematically scan 226 samples for the presence of eukaryotic sequences. Six relatively complete fungal genomes were recovered from fecal samples from three infants (Figure 3). Three are *Candida albicans*, and were reconstructed from two different infants. The two genomes from the same infant are indistinguishable, and very closely related to that from the third infant. All three are closely related to, but distinguishable from the *C*. *albicans* reference strain WO-1 (Figure 3, **Supplementary Figure S4A**). The other three fungal genomes are strains of *Candida parapsilosis* that all occurred in a single infant. These are essentially indistinguishable from each other and from the C. *parapsilosis* strain CDC317 reference genome, with which they share >99.7% average nucleotide identity (ANI) (Figure 3, **Supplementary Figure S4A,B**) (Sharon et al. 2013, Raveh-Sadka et al. 2015, Raveh-Sadka et al. 2016). *C*. *albicans* and *C*. *parapsilosis* are both clinically relevant human pathogens (Trofa et al. 2008, Kim et al. 2011).

Within thiocyanate reactor samples, genomes of a Rotifer, Rhizaria and a relative of the slime mold *Fonticula alba* had previously been identified (Kantor et al. 2015, Kantor et al. 2017). With EukRep we were able to rapidly identify these eukaryotic genomes and evaluate their completeness. Genome completeness analysis benefited from improved gene predictions for single-copy orthologous genes and showed that the identified genomes ranged in completeness from 69%-91%. (Figure 3). As previously reported (Kantor et al. 2017), the Rotifer was present in seven different samples (**Rotifer_A-G**, Figure 3), consistent with its persistence in the thiocyanate reactor community. All seven bins shared greater than 99% ANI (**Supplemental Figure S4B**) indicating they are likely the same species.

## Discussion

Using a newly acquired and two previously reported whole community metagenomic datasets we demonstrated that it is possible to rapidly recover high quality eukaryotic genomes from metagenomes for phylogenetic and metabolic analyses. The key step implemented in this study was the pre-sorting of eukaryotic genome fragments prior to gene prediction. By training and using eukaryotic gene predictors we achieved much higher quality eukaryotic gene predictions than those obtained using a prokaryotic gene prediction algorithm on the entire dataset (i.e., without separation based on phylogeny). This was critical for draft genome recovery and evaluation of genome completeness.

Classification of assembled genome fragments at the Domain level was surprisingly accurate, with 98.0% (Figure 2B) of eukaryotic sequences being correctly identified as eukaryotic, despite no close relative in the training set in many cases (**Supplementary Table S2**). The high accuracy of separation suggests some underlying pattern of kmer frequencies that is different in eukaryotes compared to prokaryotes. In part, the signature may arise from different codon use patterns associated with the different genetic codes for bacteria and eukaryotes.

We anticipate that reexamination of environmental metagenomic datasets using the same approach as implemented here will yield high quality genomes for previously unknown eukaryotes. An important benefit from this and future sequencing efforts will be an expanded knowledge of the diversity, distribution and functions of microbial eukaryotes, which are widely acknowledged as understudied (Pawlowski et al. 2012). Increasing the diversity of sequenced eukaryotic genomes would benefit evolutionary studies. Current eukaryotic multigene trees form a solid backbone of the eukaryotic tree of life (Parfrey et al. 2010) but suffer from sparse eukaryotic taxon sampling. Single gene trees, which are possible to construct from gene surveys, lack the resolution of multigene trees (Rokas and Carroll 2005). Comprehensive sequencing of full genomes would help diminish the sparse taxon sampling problem in multigene trees and improve eukaryotic evolutionary reconstructions, with implications for understanding of eukaryotic protein function. For example, Ovchinnikov et al. (2017) demonstrated that it is possible to accurately predict protein structure by utilizing residue-residue contacts inferred from evolutionary data, but such analyses require large numbers of aligned sequences. More diverse eukaryotic sequences could expand the utility of this method for eukaryotic protein family analyses. Furthermore, a broader diversity of eukaryotic genomes would provide new insights regarding gene transfer patterns and whole genome evolution.

EukRep, applied in the context of metagenomics, may prove useful for genome sequencing projects where isolation of the organism of interest may be difficult or not technically feasible. For example, it could be applied to study populations of bacteria within the hyphae of arbuscular mycorrhizal fungi (Hoffman et al. 2010).

Eukaryotic cells frequently contain multiple sets of chromosomes (diploid or polyploid). These are often very similar, but not identical, and can result in the genome assembly alternating between collapsing and splitting contigs representing homologous genomic regions (Margarido et al.). If reads are only allowed to map to one location when determining genome coverage, this could lead to variation of coverage values across different portions of a genome. As differential coverage of contigs is a parameter commonly used to help bin genomes, ploidy can complicate genome recovery. Another potential problem could relate to contamination of eukaryotic genome bins with some bacterial fragments. This will occur to some extent, given that some bacterial and archaeal contigs were wrongly classified as eukaryotic. Phylogenetic profiling of the predicted genes can be used to screen out most prokaryotic sequences.

During development we noted that the frequency of correct identification of bacterial genomes was improved by increasing the number and diversity of eukaryote sequences used in classifier training. Further improvements are anticipated as the variety of reference sequences increases. However, there may be biological reasons underpinning incorrect profiles. The small number of cases where EukRep profiled bacteria as eukaryotes or vice versa may be interesting targets for further analysis. Notably, almost all are inferred or known symbionts or parasites, raising the question of whether their sequence composition has evolved to mirror that of their hosts.

We demonstrated the value of EukRep-enabled analyses through study of an ecosystem that had been perturbed by addition of a carbon source. The results clearly show a large shift in the community composition and selection for fungi. Of the binned genomes, the fungi have by far the most cellulose-, hemicellulose-, and pectin-degrading enzymes, consistent with their enrichment in response to high organic carbon availability from degrading wood. We also genomically characterized what appears to be a macroscopic hexapod that is related to springtails (Collembola), organisms known to feed on fungi (Chen et al. 1996). Given that the hexapod genome has a large number of chitin-degrading enzymes (Supplemental Table S3), we speculate that it may be part of the community supported by the fungi in the decaying wood. However, it is also possible that it was associated with the wood prior to its addition to the geyser conduit. Interestingly, the eukaryote-based community contains very few members of the candidate phyla radiation (CPR) and an archaeal radiation known as DPANN and other CP bacteria. These novel organisms are mostly predicted to be anaerobes and are highly abundant in groundwater samples that were likely sourced from deep aquifers under the Colorado Plateau (Probst et al. 2017 in revision). The results of the current study indicate that CPR and DPANN in the Crystal Geyser system are adapted to an environment with relatively low in carbon availability, a finding that may guide future laboratory enrichment studies that target these organisms.

Overall, the results reported here demonstrate that comprehensive, cultivation-independent genomic studies of ecosystems containing a wide variety of organisms types are now possible. Examples of future applications include analysis of the distribution and metabolic capacities and potential pathogenicity of fungi in the human microbiome, tracking of eukaryotes, (including multicellular eukaryotes) in reactors used in biotechnologies, profiling of the built environment and natural ecosystem research.

## Methods

### Crystal Geyser Sample Collection and DNA extraction

Filtration of groundwater for sample CG_bulk is given in Probst et al 2017 (sample CG23_combo_of_CG06-09_8_20_14). Groundwater containing particulate wood was collected in a 50-ml falcon tube. All samples were frozen on site on dry ice and stored at −80 °C until further processing. The sample with the particulate wood was spun down and DNA extraction was performed as described previously (Emerson et al, 2016).

### Crystal Geyser DNA Sequencing and Assembly

Raw sequencing reads were processed with bbtools (http://jgi.doe.gov/data-and-tools/bbtools/) and quality filtered with SICKLE with default parameters (Version 1.21, https://github.com/najoshi/sickle). IBDA_UD (Peng et al., 2012) was used to assemble and scaffold filtered reads. Scaffolding errors were corrected using MISS (Sharon et al., unpublished), a tool that searches and fixes gaps in the assembly based on mapped reads that exhibit inconsistencies between raw reads and assembly. The two Crystal Geyser samples used for binning and comparison in this study, CG_WC and CG_bulk, resulted in 874 and 529 Mbps of assembled scaffolds respectively.

### Prokaryotic Genome Binning and Annotations

Protein-coding genes were predicted on entire metagenomic samples using Prodigal (Hyatt et al., 2010). Ribosomal RNA genes were predicted with Rfam (Nawrocki et al., 2015) and 16S rRNA genes were identified using SSU-Align (Nawrocki, 2009). Predicted proteins were functionally annotated by finding the best blast hit using USEARCH (ublast, Edgar, 2010) against UniProt (UniProt, 2010), Uniref90 (Suzek et al., 2007), and KEGG (Kanehisa et al. 2016). Prokaryotic draft genomes were binned through the use of emergent self-organizing map (ESOM)-based analyses of tetranucleotide frequencies. Bins were then refined through the use of ggKbase (ggkbase.berkeley.edu) to manually check the GC, coverage, and phylogenetic profiles of each bin.

### EukRep Training and Testing

EukRep along with trained linear SVM classifiers are available at https://github.com/patrickwest/EukRep. A diverse reference set of 194 bacterial genomes, 218 archaeal genomes, 27 Opisthokonta and 43 Protist genomes was obtained from NCBI and JGI (**Supplemental Table S1**). The contigs comprising these genomes were split into 5 kb chunks for which 5-mer frequencies were calculated. Contigs shorter than 3 kb were excluded. The 5-mer frequencies were used to train a linear-SVM (scikit-learn, v. 0.18, default parameters with C=100) to classify sequences as either of Opisthokonta, Protist, bacterial, or archaeal origin. The hyper-parameter C was optimized using a grid-search with cross-validation and accuracy on a subset of test genomes used for scoring. To classify an unknown or test sequence, the sequence was split into 5 kb chunks and 5-mer frequencies were determined for each chunk. Contigs shorter than 3 kb were excluded. The trained classifier was then used to predict whether the sequence is of Opisthokonta, Protist, bacterial, or archaeal origin. Once classified, the 5kb chunks were stitched back together into their parent contig and the parent contig’s taxonomy was determined based upon majority rule of its 5 kb. Accuracy for a given genome was considered to be the percent of total base pairs correctly identified as either eukaryotic or prokaryotic. To compare the effect of kmer length on prediction accuracy, kmer frequencies ranging in length from 4-6bp from the same training set were used to train separate linear-SVM models. To determine the minimum sequence length cutoff, test genomes were fragmented into pieces of n length and sequences shorter than n length were filtered out.

To test EukRep, a separate set of 97 eukaryotic and 393 prokaryotic genomes was obtained from NCBI and JGI (**Supplemental Table S2**). Genomes assembled into less than 10 contigs were fragmented into 100 kb pieces in order to better represent metagenomic datasets. EukRep was then run on each genome individually. Accuracy for a given genome was measured by dividing the total number of bps correctly classified by the total number of bps tested.

### Eukaryotic Genome Binning and Annotations

Scaffolds predicted to be eukaryotic scaffolds by EukRep were binned into putative genomes using CONCOCT (Alneberg et al. 2014). Eukaryotic genome bins smaller than 5 Mbp were not included in further analyses. Gene predictions were performed individually on each bin with the MAKER2 pipeline (Holt and Yandell, 2011, v. 2.31.9) with default parameters and using GeneMark-ES (Ter-Hovhannisyan et al. 2008, v. 4.32), AUGUSTUS (Stanke et al. 2006, v. 2.5.5) trained with BUSCO (Simão et al. 2015, v. 2.0), and the proteomes of *C*. *Reinhardtti* (Merchant et al. 2007), *N*. *Crassa* (Galagan et al. 2003), and *R*. *filosa* (Glöckner et al. 2014) for homology evidence. These gene prediction strategies were employed due to their ability to be automatically trained for individual genomes. Completeness of the combined MAKER predicted gene set as well as the individual gene predictor gene sets were compared and the most complete based upon BUSCO analysis was used in future analyses. Phylogenetic classification of the predicted genes along with presence or absence of single-copy orthologous genes was then used to refine each binned genome. CAZYmes were detected in both eukaryotic and prokaryotic bins through the use of HMMER3 (Eddy 1998, v. 3.1b2) and a set of HMMs obtained from dbcan (Yin et al. 2012). The presence or absence of various metabolic pathways was determined by using a custom set of metabolic pathway marker gene HMMs (Anantharaman et al. 2016) and HMMER3. Protease and lipases were predicted by using lipase HMMs from the Lipase Engineering Database (Fischer and Pleiss 2003), and blasting against a protease database obtained from MEROPS (Rawlings et al. 2016). Putative excreted proteases and lipases were identified by searching for predicted proteases and lipases with secretion signals identified with SignalP (Petersen et al. 2011) and no more than one transmembrane domain with TMHMM (Krogh et al, 2001). To find potentially contaminating prokaryotic scaffolds, predicted genes were blasted against Uniprot. Scaffolds in which the majority of best hits belonged to prokaryotic genes were removed.

### Eukaryotic Gene Set Comparisons

Nine gene sets were obtained from JGI’s mycocosm database (Grigoriev et al. 2011) and NCBI. For each genome, genes were predicted without transcriptomic evidence by running assembled sequences through the MAKER pipeline with AUGUSTUS trained with BUSCO, and GeneMark-ES in self-training mode. Gene sets predicted with transcriptomic evidence were obtained from the JGI portal and NCBI. For comparison against eukaroytic Prodigal predicted gene sets, Prodigal was run with the ‘-meta’ flag.

### Eukaryote Genome Completeness Estimates

Genome completeness of predicted eukaryotic genomes was estimated based on the presence of conserved, low-copy-number genes. BUSCO (Simão et al. 2015, v. 2.0) was run with default parameters using the “eukaryota_odb9” lineage set composed of 303 core eukaryotic genes. Completeness was considered to be the percent of the total 303 core genes that were present in either single or duplicated copies. Additionally, the number of genes identified as duplicated was used as a way to estimate how much of a given binned genome appeared to be from a single organism.

### Phylogenetic Analyses

To determine ANI between genomes, dRep was used (Olm et al. 2017). To estimate taxonomic composition of Crystal Geyser samples, rpS3 proteins were searched against KEGG (Kanehisa et al. 2016) with USEARCH (ublast, Edgar, 2010) and the taxonomy of the top hit was used to assign identified rpS3s to taxonomic groups. Abundance of identified rpS3s was determined by calculating the average coverage depth of the scaffolds containing annotated ribosomal protein S3 (rpS3) genes. Average coverage depth was calculated by dividing the number of reads mapped to the scaffold by the scaffold length. Abundances were normalized for comparison across samples by multiplying the average coverage depth by the sample read count times read length. 461 protein sets were obtained from binned eukaryotic genomes, publicly available genomes from the Joint Genome Institute’s IMG-M database (Chen et al. 2016, img.jgi.doe.gov), NCBI, the Candida Genome Database (Skrzypek et al.), and a previously developed data set (Hug et al. 2016). For each protein set, 16 ribosomal proteins (L2, L3, L4, L5, L6, L14, L16, L18, L22, L24, S3, S8, S10, S17 and S19) were identified by taking BLASTing a reference set of 16 ribosomal proteins obtained from a variety of protistan organisms against the protein sets. Blast hits were filtered to a minimum e-value of 1.0e^−5^ and minimum target coverage of 25%. The 16 ribosomal protein datasets were aligned with MUSCLE (v, Edgar 2004), and trimmed by removing columns containing 90% or greater gaps. The alignments were then concatenated. A maximum likelihood tree was constructed using RAxML v. 8.2.10 (Stamatakis 2014), on the CIPRES web server (Miller et al. 2010), with the LG plus gamma model of evolution (PROTGAMMALG), and with the number of bootstraps automatically determined with the MRE-based bootstopping criterion.

## Data Access

EukRep along with trained linear SVM classifiers are available at https://github.com/patrickwest/EukRep. The read datasets along with the newly assembled and binned genomes are in the process of being submitted to the NCBI Sequence Read Archive (SRA; http://www.ncbi.nlm.nih.gov/sra) and will be publically available by the time of publication. Read datasets for previously published metagenomes are available under SRA accession numbers SRA052203 and SRP056932, and BioProjects PRJNA294605 and PRJNA279279.

## Acknowledgements

This material is based upon work supported by the National Science Foundation Graduate Research Fellowship under Grant No. DGE 1106400. This study was partially funded by the Sloan Foundation (“Deep Life”, grant no. G-2016-20166041). We thank MR Olm and Dr. CT Brown for their contributions to this study.

